## Supplementary Table 1 for "A wake-active locomotion circuit depolarizes a sleep-active neuron to switch on sleep"

**S1 Table. Primers used in this study.**

Primer sequence 5’-3’

*aptf-1(gk794)*

CGACAATCTTCCCAAAGACC

CGGATCGATTGCTAGAGAGG

GCTTGGACGGCTTTAGTTGA

ArchT

ACTTCATCGTCAAGGGATGG

CATGCAGATGGTGGAGAAGA

*eat-4(ky5)*

GGGGCGTTTCCTTTTCTTTA

AAAATGCTCCGACTCCGATT

ACAGATCCATACGGAAAAGTTC

*flp-11(tm2706)*

CAGGAGTTGTTCGAGCAGAA

TCGTCCAATGGAGACCTCTT

TAGCCGCTCGTCTCACTTTT

*flp-18(db99)*

CGAACGAATCAGCCATGTAA

GAGATTCGACGATGACACGA

GGCTTGGGAGGAAGATTTTT

ICE

CCGAGCTTTGATTGACTCCG

AGTCATGTCCGAAGCAGTGA

*nmr-1(ak4)*

TGCTGGTGACTTATGAGCCT

TGCTGGCGATCTTACTGGAA

CAACACCGATGCAGAGCTC

*tdc-1(n3420)*

GAGGATCCACGCCAGAATGA

CATGTGAATCCGCCCAGAAG
