## Supplementary Table 2 for "A wake-active locomotion circuit depolarizes a sleep-active neuron to switch on sleep"

**S2 Table. Plasmids used for this study.**

| **plasmid name** | **transgene** | **length of transgene (kb)** | **source / reference** |
| --- | --- | --- | --- |
| pMK-RQ-aptf-1p | *aptf-1p* | 1,500 | [1] |
| pTNZ001_pENTR_4-1_dat1p | *dat-1p* | 784 | [2] |
| pMK-RQ-flp-11p | *flp-11p* | 2,746 | [3] |
| pPUC57-gcy-13p | *gcy-13p* | 2,192 | [4] |
| pPUC57-kan-tbh-1p | *tbh-1p* | 1,407 | [5] |
| pKA1261 hlh-34 prom | *hlh-34p* | 2,500 | [6] |
| pEntr P4-P1R plad-2 | *lad-2p* | 4,038 | [7] |
| pEntrL4-R1 pnmr-1 (pIR11) | *nmr-1p* | 2,244 | [8] |
| pEntrL4-R1 psra-6 | *sra-6p* | 4,090 | [9] |
| pTNZ024-5_pENTR_4-1_tdc-1p | *tdc-1p* | 3,452 | [5, 10] |
| pMK-tol-1p | *tol-1p* | 3,988 | [11] |
| pBSK(+)-Kan-zk673.11p | *zk673.11p* | 2,000 | [12] |
| pMK-SL1GCaMP3.35-SL2 | *GCaMP3.35* | 1,865 | [13] |
| pMK-RQ-ArchT | *ArchT* | 955 | [14] |
| pMK-ReaChR-STOP | *ReaChR-STOP* | 1,412 | [15] |
| pMK-egl-1 | *egl-1* | 376 | [14] |
| pMK-RQ-SL2mKate2unc-54-3UTR | *mKate2::unc-54 3’ UTR* | 1,995 | [13] |

**References**

1.	Turek M, Lewandrowski I, Bringmann H. An AP2 transcription factor is required for a sleep-active neuron to induce sleep-like quiescence in C. elegans. Curr Biol. 2013;23(22):2215-23. Epub 2013/11/05. doi: 10.1016/j.cub.2013.09.028. PubMed PMID: 24184105.

2.	Ezcurra M, Tanizawa Y, Swoboda P, Schafer WR. Food sensitizes C. elegans avoidance behaviours through acute dopamine signalling. EMBO J. 2011;30(6):1110-22. Epub 2011/02/10. doi: emboj201122 [pii]

10.1038/emboj.2011.22. PubMed PMID: 21304491.

3.	Turek M, Besseling J, Spies JP, Konig S, Bringmann H. Sleep-active neuron specification and sleep induction require FLP-11 neuropeptides to systemically induce sleep. eLife. 2016;5. Epub 2016/03/08. doi: 10.7554/eLife.12499. PubMed PMID: 26949257; PubMed Central PMCID: PMC4805538.

4.	Ortiz CO, Etchberger JF, Posy SL, Frokjaer-Jensen C, Lockery S, Honig B, et al. Searching for neuronal left/right asymmetry: genomewide analysis of nematode receptor-type guanylyl cyclases. Genetics. 2006;173(1):131-49. Epub 2006/03/21. doi: 10.1534/genetics.106.055749. PubMed PMID: 16547101; PubMed Central PMCID: PMCPMC1461427.

5.	Alkema MJ, Hunter-Ensor M, Ringstad N, Horvitz HR. Tyramine Functions independently of octopamine in the Caenorhabditis elegans nervous system. Neuron. 2005;46(2):247-60. Epub 2005/04/26. doi: S0896-6273(05)00167-4 [pii]

10.1016/j.neuron.2005.02.024. PubMed PMID: 15848803.

6.	Cunningham KA, Hua Z, Srinivasan S, Liu J, Lee BH, Edwards RH, et al. AMP-activated kinase links serotonergic signaling to glutamate release for regulation of feeding behavior in C. elegans. Cell Metab. 2012;16(1):113-21. Epub 2012/07/10. doi: 10.1016/j.cmet.2012.05.014. PubMed PMID: 22768843; PubMed Central PMCID: PMCPMC3413480.

7.	Wang X, Zhang W, Cheever T, Schwarz V, Opperman K, Hutter H, et al. The C. elegans L1CAM homologue LAD-2 functions as a coreceptor in MAB-20/Sema2 mediated axon guidance. J Cell Biol. 2008;180(1):233-46. Epub 2008/01/16. doi: 10.1083/jcb.200704178. PubMed PMID: 18195110; PubMed Central PMCID: PMCPMC2213605.

8.	Ben Arous J, Tanizawa Y, Rabinowitch I, Chatenay D, Schafer WR. Automated imaging of neuronal activity in freely behaving Caenorhabditis elegans. J Neurosci Methods. 2010;187(2):229-34. Epub 2010/01/26. doi: 10.1016/j.jneumeth.2010.01.011. PubMed PMID: 20096306.

9.	Hilliard MA, Apicella AJ, Kerr R, Suzuki H, Bazzicalupo P, Schafer WR. In vivo imaging of C. elegans ASH neurons: cellular response and adaptation to chemical repellents. EMBO J. 2005;24(1):63-72. Epub 2004/12/04. doi: 7600493 [pii]

10.1038/sj.emboj.7600493. PubMed PMID: 15577941.

10.	Guo ZV, Hart AC, Ramanathan S. Optical interrogation of neural circuits in Caenorhabditis elegans. Nat Methods. 2009;6(12):891-6. Epub 2009/11/10. doi: nmeth.1397 [pii]

10.1038/nmeth.1397. PubMed PMID: 19898486.

11.	Brandt JP, Ringstad N. Toll-like Receptor Signaling Promotes Development and Function of Sensory Neurons Required for a C. elegans Pathogen-Avoidance Behavior. Curr Biol. 2015;25(17):2228-37. Epub 2015/08/19. doi: 10.1016/j.cub.2015.07.037. PubMed PMID: 26279230; PubMed Central PMCID: PMCPMC4642686.

12.	Gonzales DL, Zhou J, Fan B, Robinson JT. Microfluidic-Induced Sleep: A Spontaneous C. elegans Sleep State Regulated by Satiety, Thermosensation and Mechanosensation. BioArxive. 2019. doi: <https://doi.org/10.1101/547075>

13.	Schwarz J, Lewandrowski I, Bringmann H. Reduced activity of a sensory neuron during a sleep-like state in Caenorhabditis elegans. Curr Biol. 2011;21(24):R983-4. Epub 2011/12/24. doi: S0960-9822(11)01207-3[pii]10.1016/j.cub.2011.10.046. PubMed PMID: 22192827.

14.	Wu Y, Masurat F, Preis J, Bringmann H. Sleep Counteracts Aging Phenotypes to Survive Starvation-Induced Developmental Arrest in C. elegans. Curr Biol. 2018;28(22):3610-24 e8. Epub 2018/11/13. doi: 10.1016/j.cub.2018.10.009. PubMed PMID: 30416057; PubMed Central PMCID: PMCPMC6264389.

15.	Urmersbach B, Besseling J, Spies JP, Bringmann H. Automated analysis of sleep control via a single neuron active at sleep onset in C. elegans. Genesis. 2016;54(4):212-9. Epub 2016/02/03. doi: 10.1002/dvg.22924. PubMed PMID: 26833569.
