## Supplementary Text 1 for "A wake-active locomotion circuit depolarizes a sleep-active neuron to switch on sleep"

**S1 Text. Strain list.**

The following *C. elegans* strains were used for this study:

HBR227	*aptf-1(gk794) II.* [1]

HBR430	*goeIs64 [aptf-1p::SL1-GCaMP3.35-SL2::mKate2-unc-54-3’utr, unc-119(+)].* [1]

HBR448	*aptf-1(gk794) II; goeIs64 [aptf-1p::SL1-GCaMP3.35-SL2::mKate2-unc-54-3’utr, unc-119(+)].* [1]

HBR531	*yxIs1 [glr-1p::GCaMP3, unc-122p::gfp]; aptf-1(gk794) II.* [1]

HBR560	*goeIs120 [tdc-1p::SL1-GCaMP3.35-SL2::mKate2-unc-54-3’utr, unc119(+)].* Generated for this study.

HBR1009	*flp-11(tm2706)X, goeIs118 [aptf-1p::SL1-GCaMP3.35-SL2::mKate2-aptf-1-3’utr, unc-119(+)].* [2]

HBR1118	*aptf-1(gk794) II; goeIs120 [tdc-1p::SL1-GCaMP3.35-SL2::mKate2-unc-54-3’utr, unc119(+)].* Generated for this study.

HBR1228	*goeIs268 [aptf-1p::SL1-GCaMP3.35-SL2::aptf-1-3’utr, unc-119(+)].* [1]

HBR1361	*goeIs304 [flp-11p::SL1-GCaMP3.35-SL2::mKate2-unc-54-3’UTR, unc-119(+)].* [3]

HBR1374	*goeIs307 [flp-11p::ArchT::SL2mKate2-unc-54-3’utr,unc-119(+)]; goeIs304 [flp-11p::SL1-GCaMP3.35-SL2::mKate2-unc-54-3’UTR, unc-119(+)].* [3]

HBR1391	*goeIs268 [aptf-1p::SL1-GCaMP3.35-SL2::aptf-1-3'utr, unc-119(+)]; goeIs273 [tdc-1p::ReaChR::mKate2-unc-54-3'utr, unc-119(+)].* Generated for this study.

HBR1394	*goeIs268 [aptf-1p::SL1-GCaMP3.35-SL2::aptf-1-3'utr, unc-119(+)]; goeIs273 [tdc-1p::ReaChR::mKate2-unc-54-3'utr, unc-119(+)]; tdc-1 (n3420) II.* Generated for this study, *tdc-1(n3420)* was crossed from MT10661 [4-6].

HBR1465	*goeIs120 [tdc-1p::SL1-GCaMP3.35-SL2::mKate2-unc-54-3'utr, unc119(+)]; goeIs315 [flp-11p::ReaChR::mKate2-unc-54-3'UTR, unc-119(+)].* Generated for this study.

HBR1466	*goeIs268 [aptf-1p::SL1-GCaMP3.35-SL2::aptf-1-3’utr, unc-119(+)]; goeIs315 [flp-11p::ReaChR::mKate2-unc-54-3’UTR, unc-119(+)].* Generated for this study.

HBR1472	*goeIs268 [aptf-1p::SL1-GCaMP3.35-SL2::aptf-1-3’utr unc-119(+)]; goeIs307 [flp-11p::ArchT::SL2mKate-2-unc-54-3’utr, unc-119(+)].* Generated for this study.

HBR1478	*goeIs268 [aptf-1p::SL1-GCaMP3.35-SL2::aptf-1-3’utr, unc-119(+)]; goeEx557 [gcy-13p::ArchT::mKate-2-unc-54-3´-utr, unc-119(+)]*. Generated for this study.

HBR1482	*goeIs268 [aptf-1p::SL1-GCaMP3.35-SL2::aptf-1-3’utr, unc-119(+)]; goeEx561 [gcy-13p::ReaChR-mKate-2-unc-54-3´-utr, unc-119(+)]*. Generated for this study.

HBR1533	*goeIs268 [aptf-1p::SL1-GCaMP3.35-SL2::aptf-1-3'utr, unc-119(+)]; goeIs273 [tdc-1p::ReaChR::mKate2-unc-54-3'utr, unc-119(+)]; flp-18(db99) X.* Generated for this study, *flp-18(db99)* was crossed from AX1410 [4,5,7].

HBR1537	*goeIs268 [aptf-1p::SL1-GCaMP3.35-SL2::aptf-1-3’utr, unc-119(+)]; goeIs308 [dat-1p::ReaChR::mKate2-unc-54-3’UTR, unc-119(+)].* Generated for this study.

HBR1572	*goeIs268 [aptf-1p::SL1-GCaMP3.35-SL2::aptf-1-3'utr, unc-119(+)]; goeIs273 [tdc-1p::ReaChR::mKate2-unc-54-3'utr, unc-119(+)]; flp-18(db99) X; tdc-1(n3420) II.* Generated for this study, *flp-18(db99)* was crossed from AX1410 [4,5,7] *and tdc-1(n3420) was crossed from MT10661* [6].

HBR1589	*goeIs268 [aptf-1p::SL1-GCaMP3.35-SL2::aptf-1-3’utr, unc-119(+)]; goeIs330 [nmr-1p::ArchT::mKate-2-unc-54-3’utr, unc-119(+)].* Generated for this study.

HBR1597	*goeIs268 [aptf-1p::SL1-GCaMP3.35-SL2::aptf-1-3’utr, unc-119(+)], goeIs332 [nmr-1p::ReaChR::mKate2-unc-54-3’utr, unc-119(+)].* Generated for this study.

HBR1659	*unc-119(ed3) III; goeIs364 [tdc-1p::egl-1::SL2-mKate2-unc-54-3’utr, unc-119(+)].* Generated for this study.

*HBR1753	wtfIs5 [rab-3p::NLS::GCaMP6s; rab-3p::NLS::tagRFP].* Generated for this study, *wtfIs5* was backcrossed from AML32 with N2 twice [8].

HBR1776	*wtfIs5 [rab-3p::NLS::GCaMP6s; rab-3p::NLS::tagRFP]; goeIs307 [flp-11 p::ArchT::SL2mKate2-unc-54-3’utr, unc-119(+)].* Generated for this study, *wtfIs5* was crossed from AML32 [4,5,8].

HBR1793	*goeIs268 [aptf-1p::SL1-GCaMP3.35-SL2::aptf-1-3’utr, unc-119(+)]; goeIs293 [tol-1p::ReaChR::mKate2-unc-54-3’utr, unc-119(+)].* Generated for this study.

HBR1807	*goeIs232 [sra-6p::ReaChR::mKate2-unc-54-3’utr, unc-119(+)]; goeIs304 [flp-11p::SL1-GCaMP3.35-SL2::mKate2-unc-54-3’UTR, unc-119(+)].* [3]

HBR1844	*goeIs268 [paptf-1p::SL1-GCaMP3.35-SL2::aptf-1-3’utr, unc-119(+)]; goeIs340 [dat-1 p::ArchT::SL2mKate2-unc-54-3UTR].* Generated for this study.

HBR1845	*goeIs268 [paptf-1p::SL1-GCaMP3.35-SL2::aptf-1-3’utr, unc-119(+)]; goeIs370 [lad-2p::ReaChR::mKate2-unc-54-3’UTR, unc-119(+)].* Generated for this study.

HBR1849	*goeIs304 [flp-11p::SL1-GCaMP3.35-SL2::mKate2-unc-54-3’UTR,*

*unc-119(+)]; goeIs364 [tdc-1p::egl-1::SL2-mkate2-unc-54-3’utr,*

*unc-119(+)]; goeIs232 [sra-6p::ReaChR::mKate2-unc-54-3’utr,*

*unc-119(+)].* Generated for this study.

HBR1873	*goeIs268 [paptf-1p::SL1-GCaMP3.35-SL2::aptf-1-3’utr, unc-119(+)]; goeIs373 [lad-2p::ArchT::SL2-mKate2-unc-54-3’UTR, unc-119(+)].* Generated for this study.

HBR1889	*goeIs120 [tdc-1p::SL1-GCaMP3.35-SL2::mKate2-unc-54-3’utr, unc119(+)]; goeIs232 [sra-6p::ReaChR::mKate2-unc-54-3’utr, unc-119(+)].* Generated for this study.

HBR1951	*ynIs40 [flp-11p::GFP] V; goeIs359 [nmr-1p::egl-1::SL2-mkate2-unc-54-3’utr, unc-119(+)].* Generated for this study, *ynIs40 [flp-11p::GFP]* was crossed from NY2040 [9].

HBR1952	*goeIs304 [flp-11p::SL1-GCaMP3.35-SL2::mKate2-unc-54-3'UTR, unc-119(+)]; goeIs343 [tdc-1p::ArchT::mKate2-unc-54-3'utr, unc-119(+)].* Generated for this study.

HBR1967	*goeIs304 [flp-11p::SL1-GCaMP3.35-SL2::mKate2-unc-54-3'UTR, unc-119(+)], goeEx705 [nmr-1p::ArchT::mKate-2-unc-54-3'utr, unc-119(+)].* Generated for this study.

HBR1982	*goeIs402 [tol-1p::ArchT::SL2-mKate2-unc-54-3’UTR,unc-119(+)]; goeIs304 [flp-11p::SL1-GCaMP3.35-SL2::mKate2-unc-54-3’UTR, unc-119(+)].* Generated for this study.

HBR2019	*akIs11 [nmr-1p::ICE]; goeIs307 [flp-11p::ArchT::SL2mKate2-unc-54-3’utr, unc-119(+)]; goeIs304 [flp-11p::SL1-GCaMP3.35-SL2::mKate2-unc-54-3’UTR, unc-119(+)].* Generated for this study, *akIs11* was obtained from A. V. Maricq [10].

HBR2021	*goeIs307 [flp-11p::ArchT::SL2mKate2-unc-54-3’utr,unc-119(+)]; goeIs304 [flp-11p::SL1-GCaMP3.35-SL2::mKate2-unc-54-3’UTR, unc-119(+)]; nmr-1(ak4) II.* Generated for this study, *ak4* was crossed from VM487 [11].

HBR2033	*goeIs195 [nmr-1p::SL1-GCaMP6s::mKate2-unc-54-3’utr, unc-119(+)]; goeIs403 [flp-11p::ArchT::mKate2-flp-11-3’utr, unc-119(+)].* Generated for this study.

HBR2039	*goeIs307 [flp-11p::ArchT::SL2mKate2-unc-54-3'utr, unc-119(+)]; goeIs120 [tdc-1p::SL1-GCaMP3.35-SL2::mKate2-unc-54-3'utr, unc119(+)].* Generated for this study.

HBR2058	*goeIs304 [flp-11p::SL1-GCaMP3.35-SL2::mKate2-unc-54-3'UTR, unc-119(+)]; goeEx716 [tbh-1p::ReaChR::mKate2unc-54 3'UTR, unc119(+); unc-122::RFP].* Generated for this study.

HBR2109	*goeIs195 [nmr-1p::SL1-GCaMP6s::mKate2-unc-54-3’utr, unc-119(+)]; goeIs315 [flp-11p::ReaChR::mKate2-unc-54-3’UTR, unc-119(+)].* Generated for this study.

HBR2123	*goeIs268 [aptf-1p::SL1-GCaMP3.35-SL2::aptf-1-3'utr, unc-119(+)]; goeEx561 [gcy-13p::ReaChR-mkate-2-unc-54-3´-utr, unc-119(+)]; eat-4(ky5) III.* Generated for this study, *eat-4(ky5)* was crossed from MT6308 [12].

HBR2128	*goeIs304 [flp-11p::SL1-GCaMP3.35-SL2::mKate2-unc-54-3’UTR, unc-119(+)]; eat-4(ky5) III.* Generated for this study, *eat-4(ky5)* was crossed from MT6308 [12].

HBR2169	*goeEx718 [hlh-34p::ReaChR::mKate2-unc-54-3’UTR,unc-119(+); myo-2p::mCherry]; goeIs304 [flp-11p::SL1-GCaMP3.35-SL2::mKate2-unc-54-3’UTR, unc-119(+)].* Generated for this study.

HBR2180	*goeEx725 [hlh-34p::ArchT::SL2mKate2-unc-54-3’UTR, unc-119(+); myo-3p::mCherry]; goeIs304 [flp-11p::SL1-GCaMP3.35-SL2::mKate2-unc-54-3’UTR, unc-119(+)].* Generated for this study.

HBR2231	*goeIs445 [zk673.11p::ArchT::SL2mKate2-unc-54 3‘ UTR, unc-119(+)].* Generated for this study.

HBR2243	*goeIs445 [zk673.11p::ArchT::SL2mKate2-unc-54 3‘ UTR, unc-119(+)]; goeIs5 [nmr-1p::SL1-GCaMP3.35-SL2::unc-54-3‘utr, unc-119(+)].* Generated for this study.

HBR2271	*flp-11(syb816 [SL2::mKate2::linker(GSGSG)::tetanustoxin_LC])X; goeIs307 [flp-11p::ArchT::SL2mKate2-unc-54-3‘utr, unc-119(+)], goeIs304 [flp-11p::SL1-GCaMP3.35-SL2::mKate2-unc-54-3‘UTR, unc-119(+)].* Generated for this study.

HBR2272	*goeIs120 [tdc-1p::SL1-GCaMP3.35-SL2::mKate2-unc-54-3‘utr, unc-119(+)]; goeEx557 [gcy-13p::ArchT-mkate-2-unc-54-3´-utr, unc-119(+)].* Generated for this study.

HBR2274	*goeIs304 [flp-11p::SL1-GCaMP3.35-SL2::mKate2-unc-54-3‘UTR, unc-119(+)], goeIs195 [nmr-1p::SL1-GCaMP6s::mKate2-unc-54-3‘utr, unc-119(+)].* Generated for this study.

HBR2275	*goeIs307 [flp-11p::ArchT::SL2mKate2-unc-54-3‘utr,unc-119(+)]; goeIs304 [flp-11p::SL1-GCaMP3.35-SL2::mKate2-unc-54-3‘UTR, unc-119(+)]; unc-13(s69) I.* Generated for this study, *unc-13(s69)* was crossed from EG9631 [4,5].

HBR2287	*goeIs195 [nmr-1p::SL1-GCaMP6s::mkate2-unc-54-3‘utr, unc-119(+)]; goeIs332 [nmr-1p::ReaChR::mKate2-unc-54-3’utr, unc-119(+)].* Generated for this study.

HBR2288	*nsEx2846(pept-3p::TeTX, elt-2p::mCherry), goeIs195 [nmr-1p::SL1-GCaMP6s::mKate2-unc-54-3‘utr, unc-119(+)]; goeIs315 [flp-11p::ReaChR::mKate2-unc-54-3‘UTR, unc-119(+)].* Generated for this study, *pept-3p::TeTX* was crossed from OS4976 [13].

HBR2289	*goeIs195 [nmr-1p::SL1-GCaMP6s::mKate2-unc-54-3‘utr, unc-119(+)]; goeIs315 [flp-11p::ReaChR::mKate2-unc-54-3‘UTR, unc-119(+)], flp-11(tm2706)X.* Generated for this study.

HBR2316	*goeIs332 [nmr-1p::ReaChR::mKate2-unc-54-3'utr, unc-119(+)]; mzmEx324(sra-11p::mCherry, sra-11p::GCaMP5K).* Generated for this study, *sra-11p::GCaMP5K* was crossed from ZIM498 [14].

HBR2321	*goeIs332 [nmr-1p::ReaChR::mKate2-unc-54-3'utr, unc-119(+)], lite-1(ce314) X.* Generated for this study, *lite-1(ce314)* was crossed from ZIM1048 [4,5,15,16].

HBR2323	*goeIs304 [flp-11p::SL1-GCaMP3.35-SL2::mKate2-unc-54-3'UTR, unc-119(+)]; goeIs340 [dat-1p::ArchT::SL2mKate2unc-54-3’UTR]; goeIs402 [tol-1p::ArchT::SL2-mKate2-unc-54-3'UTR, unc-119(+)].* Generated for this study.

HBR2336	*goeIs120 [ptdc-1p::SL1-GCaMP3.35-SL2::mKate2-unc-54-3'utr, unc-119(+)]; goeIs273 [tdc-1p::ReaChR::mKate2-unc-54-3'utr, unc-119(+)].* Generated for this study.

The following strains not created in the lab were used:

AML32	*wtfIs5 [Prab-3::NLS::GCaMP6s; Prab-3::NLS::tagRFP].* [8]

CX14845	*kyEx4863 [rig-3p::HisCl1:sl2mCherry].* [17]

N2	Wild type (Bristol) [18]

OS4976	*nsEx2846 (pept-3p::TeTX, elt-2p::mCherry).* [13]

PHX816	*flp-11(syb816 [SL2::mKate2::linker(GSGSG)::tetanustoxin_LC]). X.*

Generated by SunyBiotech according to our design for this study.

ZC1148	*yxIs1 [glr-1p::GCaMP3.35, unc-122p::gfp].* [19]

ZIM498	*mzmEx324(sra-11p::mCherry, sra-11p::GCaMP5K).* [14]

**References**

1.	Turek M, Lewandrowski I, Bringmann H. An AP2 transcription factor is required for a sleep-active neuron to induce sleep-like quiescence in C. elegans. Curr Biol. 2013;23(22):2215-23. Epub 2013/11/05. doi: 10.1016/j.cub.2013.09.028. PubMed PMID: 24184105.

2.	Turek M, Besseling J, Spies JP, Konig S, Bringmann H. Sleep-active neuron specification and sleep induction require FLP-11 neuropeptides to systemically induce sleep. eLife. 2016;5. Epub 2016/03/08. doi: 10.7554/eLife.12499. PubMed PMID: 26949257; PubMed Central PMCID: PMC4805538.

3.	Wu Y, Masurat F, Preis J, Bringmann H. Sleep Counteracts Aging Phenotypes to Survive Starvation-Induced Developmental Arrest in C. elegans. Curr Biol. 2018;28(22):3610-24 e8. Epub 2018/11/13. doi: 10.1016/j.cub.2018.10.009. PubMed PMID: 30416057; PubMed Central PMCID: PMCPMC6264389.

4.	Rose AM, Baillie DL. Genetic organization of the region around UNC-15 (I), a gene affecting paramyosin in Caenorhabditis elegans. Genetics. 1980;96(3):639-48. Epub 1980/11/01. PubMed PMID: 7262541; PubMed Central PMCID: PMCPMC1214366.

5.	Richmond JE, Davis WS, Jorgensen EM. UNC-13 is required for synaptic vesicle fusion in C. elegans. Nat Neurosci. 1999;2(11):959-64. Epub 1999/10/20. doi: 10.1038/14755. PubMed PMID: 10526333; PubMed Central PMCID: PMCPMC2585767.

6.	Alkema MJ, Hunter-Ensor M, Ringstad N, Horvitz HR. Tyramine Functions independently of octopamine in the Caenorhabditis elegans nervous system. Neuron. 2005;46(2):247-60. Epub 2005/04/26. doi: S0896-6273(05)00167-4 [pii]

10.1016/j.neuron.2005.02.024. PubMed PMID: 15848803.

7.	Cohen M, Reale V, Olofsson B, Knights A, Evans P, de Bono M. Coordinated regulation of foraging and metabolism in C. elegans by RFamide neuropeptide signaling. Cell Metab. 2009;9(4):375-85. Epub 2009/04/10. doi: 10.1016/j.cmet.2009.02.003. PubMed PMID: 19356718.

8.	Nguyen JP, Linder AN, Plummer GS, Shaevitz JW, Leifer AM. Automatically tracking neurons in a moving and deforming brain. PLoS computational biology. 2017;13(5):e1005517. Epub 2017/05/26. doi: 10.1371/journal.pcbi.1005517. PubMed PMID: 28545068; PubMed Central PMCID: PMC5436637.

9.	Kim K, Li C. Expression and regulation of an FMRFamide-related neuropeptide gene family in Caenorhabditis elegans. The Journal of comparative neurology. 2004;475(4):540-50. Epub 2004/07/06. doi: 10.1002/cne.20189. PubMed PMID: 15236235.

10.	Zheng Y, Brockie PJ, Mellem JE, Madsen DM, Maricq AV. Neuronal control of locomotion in C. elegans is modified by a dominant mutation in the GLR-1 ionotropic glutamate receptor. Neuron. 1999;24(2):347-61. Epub 1999/11/26. PubMed PMID: 10571229.

11.	Brockie PJ, Mellem JE, Hills T, Madsen DM, Maricq AV. The C. elegans glutamate receptor subunit NMR-1 is required for slow NMDA-activated currents that regulate reversal frequency during locomotion. Neuron. 2001;31(4):617-30. Epub 2001/09/08. PubMed PMID: 11545720.

12.	Lee RY, Sawin ER, Chalfie M, Horvitz HR, Avery L. EAT-4, a homolog of a mammalian sodium-dependent inorganic phosphate cotransporter, is necessary for glutamatergic neurotransmission in caenorhabditis elegans. J Neurosci. 1999;19(1):159-67. Epub 1998/12/31. PubMed PMID: 9870947; PubMed Central PMCID: PMCPMC3759158.

13.	Katz M, Corson F, Iwanir S, Biron D, Shaham S. Glia Modulate a Neuronal Circuit for Locomotion Suppression during Sleep in C. elegans. Cell reports. 2018;22(10):2575-83. Epub 2018/03/08. doi: 10.1016/j.celrep.2018.02.036. PubMed PMID: 29514087; PubMed Central PMCID: PMCPMC5870883.

14.	Kato S, Kaplan HS, Schrodel T, Skora S, Lindsay TH, Yemini E, et al. Global brain dynamics embed the motor command sequence of Caenorhabditis elegans. Cell. 2015;163(3):656-69. Epub 2015/10/20. doi: 10.1016/j.cell.2015.09.034. PubMed PMID: 26478179.

15.	Edwards SL, Charlie NK, Milfort MC, Brown BS, Gravlin CN, Knecht JE, et al. A novel molecular solution for ultraviolet light detection in Caenorhabditis elegans. PLoS biology. 2008;6(8):e198. Epub 2008/08/09. doi: 10.1371/journal.pbio.0060198. PubMed PMID: 18687026; PubMed Central PMCID: PMC2494560.

16.	Nichols ALA, Eichler T, Latham R, Zimmer M. A global brain state underlies C. elegans sleep behavior. Science. 2017;356(6344). Epub 2017/06/24. doi: 10.1126/science.aam6851. PubMed PMID: 28642382.

17.	Pokala N, Liu Q, Gordus A, Bargmann CI. Inducible and titratable silencing of Caenorhabditis elegans neurons in vivo with histamine-gated chloride channels. Proc Natl Acad Sci U S A. 2014;111(7):2770-5. Epub 2014/02/20. doi: 10.1073/pnas.1400615111. PubMed PMID: 24550306; PubMed Central PMCID: PMCPMC3932931.

18.	Brenner S. The genetics of Caenorhabditis elegans. Genetics. 1974;77(1):71-94. Epub 1974/05/01. PubMed PMID: 4366476.

19.	Hendricks M, Ha H, Maffey N, Zhang Y. Compartmentalized calcium dynamics in a C. elegans interneuron encode head movement. Nature. 2012;487(7405):99-103. Epub 2012/06/23. doi: 10.1038/nature11081. PubMed PMID: 22722842; PubMed Central PMCID: PMCPMC3393794.
