## Supplementary Text 2 for "A wake-active locomotion circuit depolarizes a sleep-active neuron to switch on sleep"

**S2 Text. List of constructs generated during this study.**

K31	*nmr-1p::SL1-GCaMP3.35-SL2::mKate2-unc-54-3UTR, unc-119(+)*

K78	*tdc-1p::SL1-GCaMP3.35-SL2::mKate2-unc-54-3UTR, unc-119(+)*

K133	*nmr-1p::SL1-GCaMP6s-SL2::mKate2-unc-54-3UTR, unc-119(+)*

K183	*nmr-1p::ReaChR::mKate2-unc-54-3UTR, unc-119(+)*

K189	*tdc-1p::ReaChR::mKate2-unc-54-3UTR, unc-119(+)*

K190	*tdc-1p::ArchT::mKate2 unc-54 3'UTR, unc-119(+)*

K196	*gcy-13p::ArchT::SL2 mKate2 unc-54 3'UTR, unc-119(+)*

K197	*gcy-13p::ReaChR::mKate2-unc-54-3UTR, unc-119(+)*

K200	*nmr-1p::ArchT::SL2 mKate2 unc-54 3'UTR, unc-119(+)*

K204	*tol-1p::ReaChR::mKate2-unc-54-3UTR, unc-119(+)*

K215	*flp11p::ReaChR::SL2mKate2-unc-54-3UTR, unc-119(+)*

K249	*dat-1p::ReaChR::mKate2-unc-54-3UTR, unc-119(+)*

K257	*lad-2p::ReaChR::mKate2-unc-54-3UTR, unc-119(+)*

K259	*dat-1p::ArchT::SL2 mKate2 unc-54 3'UTR, unc-119(+)*

K260	*tol-1p::ArchT::SL2 mKate2 unc-54 3'UTR, unc-119(+)*

K300	*lad-2p::ArchT::SL2 mKate2 unc-54 3'UTR, unc-119(+)*

K308	*tdc-1p::egl-1::SL2mKate2-unc-54-3UTR, unc-119(+)*

K309	*nmr-1p::egl-1::SL2mKate2-unc-54-3UTR, unc-119(+)*

K345	*tbh-1p::ReaChR::mKate2-unc-54-3UTR, unc-119(+)*

*K355	hlh-34p::ReaChR::mKate2-unc-54-3UTR, unc-119(+)*

*K356	hlh-34p::ArchT::SL2 mKate2 unc-54 3'UTR, unc-119(+)*

K364	*zk673.11p::ArchT::SL2 mKate2 unc-54 3'UTR, unc-119(+)*
