## Supplementary Text 3 for "A wake-active locomotion circuit depolarizes a sleep-active neuron to switch on sleep"

**S3 Text. Sequence of the strain PHX816.**

**PHX816: *flp-11(syb816 [SL2::mKate2::linker(GSGSG)::tetanustoxin_LC]) X.***

>flp-11b-SL2(gpd-2)-mKate2 linker (GSGSG) tetanustoxin LC
TGGCACTTCTCCTTATTGTCTTCGTTGCCGCTTCTTTTGCTCAATCTTATGATGACGTCAGgtatagttttttcttaaaacaatttttatcaattacccatataaatctattgtagTGCGGAGAAACGTGCCATGCGGAACGCCTTGGTTCGATTTGGAAGAGCTAGTGGTGGAATGAGAAATGCTCTCGTTAGATTCGGAAAGAGGTCTCCATTGGACGAGGAAGACTTTGCTCCAGAGAGCCCACTCCAGGGAAAACGGAACGGTGCCCCACAACCATTTGgtaagttgtcttaaaatttttcttccgctttttgcctttgcttcatgtgtcgtttattttgctttgcagttcgctttggccgatccggtcaactcgaccacatgcacgaccttttgtcgactcttcagAAGCTCAAGTTCGCCAACAACAAGTAATG**ACC**GAGGACGACCGTCTTCTGCTCGAACAACTCCTG**CGA**CGAATTCATCATTAAgctgtctcatcctactttcacctagttaactgcttgtcttaaaatctatgcttctctttagtatctaaaattttcctagaagcttacaagtatataaatggtctcttctcaataaaggttgtatatttattcatcttattgaatctgccatttcctcgtttttgcgagtttatataccttccaattttctttctattgtattttcaacttctaattttaattcagggaaactgcttcaacgcatcATGTCCGAGCTCATCAAGGAGAACATGCACATGAAGCTCTACATGGAGGGAACCGTCAACAACCACCACTTCAAGTGCACCTCCGAGGGAGAGGGAAAGCCATACGAGGGAACCCAAACCATGCGTATCAAGgtaagtttaaacatatatatactaactaaccctgattatttaaattttcagGCCGTCGAGGGAGGACCACTCCCATTCGCCTTCGACATCCTCGCCACCTCCTTCATGTACGGATCCAAGACCTTCATCAACCACACCCAAGGAATCCCAGACTTCTTCAAGCAATCCTTCCCAGAGGGATTCACCTGGGAGCGTGTCACCACCTACGAGGACGGAGGAGTCCTCACCGCCACCCAAGACACCTCCCTCCAAGACGGATGCCTCATCTACAACGTCAAGATCCGTGGAGTCAACTTCCCATCCAACGGACCAGTCATGCAAAAGAAGACCCTCGGATGGGAGGCCTCCACCGAGACCCTCTACCCAGCCGACGGAGGACTCGAGGGACGTGCCGACATGGCCCTCAAGCTCGTCGGAGGAGGACACCTCATCTGCAACCTCAAGgtaagtttaaacatgattttactaactaactaatctgatttaaattttcagACCACCTACCGTTCCAAGAAGCCAGCCAAGAACCTCAAGATGCCAGGAGTCTACTACGTCGACCGTCGTCTCGAGCGTATCAAGGAGGCCGACAAGGAGACCTACGTCGAGCAACACGAGGTCGCCGTCGCCCGTTACTGCGACCTCCCATCCAAGCTCGGACACCGTGGATCCGGATCCGGAATGCCAATCACCATCAACAACTTCCGTTACTCCGACCCAGTCAACAACGACACCATCATCATGATGGAGCCACCATACTGCAAGGGACTCGACATCTACTACAAGGCCTTCAAGATCACCGACCGTATCTGGATCGTCCCAGAGCGTTACGAGTTCGGAACCAAGCCAGAGGACTTCAACCCACCATCCTCCCTCATCGAGGGAGCCTCCGAGTACTACGACCCAAACTACCTCCGTACCGACTCCGACAAGGACCGTTTCCTCCAAACCATGGTCAAGCTCTTCAACCGTATCAAGAACAACGTCGCCGGAGAGGCCCTCCTCGACAAGATCATCAACGCCATCCCATACCTCGGAAACTCCTACTCCCTCCTCGACAAGTTCGACACCAACTCCAACTCCGTCTCCTTCAACCTCCTCGAGCAAGACCCATCCGGAGCCACCACCAAGTCCGCCATGCTCACCAACCTCATCATCTTCGGACCAGGACCAGTCCTCAACAAGAACGAGGTCCGTGGAATCGTCCTCCGTGTCGACAACAAGgtaagtttaaacagttcggtactaactaaccatacatatttaaattttcagAACTACTTCCCATGCCGTGACGGATTCGGATCCATCATGCAAATGGCCTTCTGCCCAGAGTACGTCCCAACCTTCGACAACGTCATCGAGAACATCACCTCCCTCACCATCGGAAAGTCCAAGTACTTCCAAGACCCAGCCCTCCTCCTCATGCACGAGCTCATCCACGTCCTCCACGGACTCTACGGAATGCAAGTCTCCTCCCACGAGATCATCCCATCCAAGCAAGAGATCTACATGCAACACACCTACCCAATCTCCGCCGAGGAGCTCTTCACCTTCGGAGGACAAGACGCCAACCTCATCTCCATCGACATCAAGAACGACCTCTACGAGAAGACCCTCAACGACTACAAGGCCATCGCCAACAAGCTCTCCCAAGTCACCTCCTGCAACGACCCAAACATCGACATCGACTCCTACAAGCAAATCTACCAACAAAAGTACCAATTCGACAAGGACTCCAACGGACAATACATCGTCAACGAGGACAAGTTCCAAATCCTCTACAACTCCATCATGTACGGATTCACCGAGATCGAGCTCGGAAAGAAGTTCAACATCAAGACCCGTCTCTCCTACTTCTCCATGAACCACGACCCAGTCAAGATCCCAAACCTCCTCGACGACACCATCTACAACGACACCGAGGGATTCAACATCGAGTCCAAGGACCTCAAGTCCGAGTACAAGGGACAAAACATGCGTGTCAACACCAACGCCTTCCGTAACGTCGACGGATCCGGACTCGTCTCCAAGCTCATCGGACTCTGCAAGAAGATCATCCCACCAACCAACATCCGTGAGAACCTCTACAACCGTACCGCCTAAaaatcatatgtttttct
